# Quantifying the mechanical properties of yeast *Candida albicans* using atomic force microscopy-based force spectroscopy

**DOI:** 10.1101/2022.10.07.511277

**Authors:** Christopher R. Jones, Zhenyu Jason Zhang, Hung-Ji Tsai

## Abstract

Fungi can adapt to a wide range of environmental stress in the wild and host milieu by employing their plastic genome and great diversity in morphology. Among different adaptive strategies, mechanical stimuli, such as changes in osmotic pressure, surface remodelling, hyphal formation, and cell divisions, could guide the physical cues into physiological responses through complex signalling network. While fungal pathogens require a pressure-driven force to expand and penetrate host tissues, quantitatively studying the biophysical properties at the host-fungal interface is critical to understand the development of fungal diseases. Microscopy-based techniques have enabled researchers to monitor the dynamic mechanics on fungal cell surface in responses to the host stress and antifungal drugs. Here, we describe a label-free, high-resolution method based on atomic force microscopy, with a step-by-step protocol to measure the physical properties in human fungal pathogen *Candida albicans*.

## 1. Introduction

Fungal pathogens can change their cell surface composition and morphology to respond to environmental stimuli and to facilitate the pathogenicity (Li and Nielsen 2017; Sudbery 2011; Gow, Latge, and Munro 2017; Chai et al. 2009). Throughout the process of cell division and hyphal expansion, a hydrostatic pressure (turgor), generated by an accumulation of solutes, including amino acids and polyols which change intracellular osmolarity, serves as a key driver for tissue invasion and colonization (Lew 2011; Hohmann 2002). Understanding the underlying mechanism and where its related physical properties act on can offer as invaluable insight into the development of fungal diseases. As such, a quantitative measurement with a nanoscopic resolution for the pressure-driven force is critical to connect the biophysical cues and physiological responses in fungi.

Microorganism cell wall provides a physical barrier to resist both internal force (turgor) (Wood Janet M. 1999; Hohmann 2002) and external perturbations (e.g., macrophage engulfment) (Erwig and Gow 2016), and its composition can constantly change in response to the cell physiological state (Gow, Latge, and Munro 2017). For example, upon heat stress, the abundance of chitin changes with a two-fold increase of surface stiffness (Pillet et al. 2014). At the host-fungal interface, innate immune cells can recognize fungal cell wall components such as β-glucan and initiate phagocytosis for fungal killing (Erwig and Gow 2016; Hopke et al. 2018). During such process, polymorphic fungal pathogens, such as *Candida albicans*, can form lengthy hyphae, where β-glucan is masked by mannan to attenuate phagosome maturation (Bain et al. 2014), and also extend their cell body to escape from phagocytic engulfment (Bain et al. 2012; Lewis et al. 2012). This complex morphogenetic program is not only governed by stress-induced signalling pathways but also a series of mechanical cues to achieve infections by overcoming host immunity (Lenardon, Munro, and Gow 2010; Brand 2012; Crampin et al. 2005; Bain et al. 2014; O’Meara et al. 2018). In parallel, macrophage can counteract the physical challenges by folding hyphae and further damaging hyphal structure while re-exposing β-glucan to promote immune recognition (Bain et al. 2021). This two-way host-fungal response highlights the important role of physical properties in fungal pathogenesis. Furthermore, to colonize host tissue or implanted abiotic surface, a sufficient adhesive force, resulting from either adhesins (e.g., Als proteins (Hoyer 2001)) or carbohydrate moiety is essential for the aforementioned pathogenic process (Gow and Hube 2012; Gulati and Nobile 2016). Additionally, fungal cell wall is a major target for current antifungal treatments, and antifungal drugs, including fluconazole and caspofungin, can alter fungal surface mechanics differently (Çolak et al. 2020; El-Kirat-Chatel et al. 2013). These surface physical properties are critical parameters for antifungal efficacy while developing strategies in fungal clearance and the measurements of antifungal drug efficacy.

Techniques employing different fluorescence probes and soft substrates to measure cell surface stiffness as a proxy of turgor changes and monitor its dynamics in response to environment (Ryder et al. 2022; Ravishankar et al. 2001; Howard et al. 1991). Over the past two decades, atomic force microscopy (AFM) has been used extensively to investigate the mechanical and topographical characteristics of living microorganisms without additional labeling or sophisticated sample preparation procedures (Zhang et al. 2010; El-Kirat-Chatel and Dufrêne 2016; Valotteau et al. 2019; Dufrêne et al. 2021). A unique advantage of this technique is to acquire nanoscopic spatial resolution under physiological condition in response to stress applied. As the method in evaluating mechanical properties in *C. albicans* is proven versatile in mycology, here we describe the measurements of cell surface stiffness and turgor pressure in *C. albicans* with step-by-step instructions.

## 2. Materials

### 2.1. Strain, medium and solutions

1. *Candida albicans* SC5314 (wild-type).
2. YPD medium (liquid and solid): 1% bacto yeast extract; 2% bacto peptone; 2% dextrose, and 1.5% bacto agar (solid media only).
3. Poly-L-Lysine (0.1% w/v, Sigma-Aldrich P8920.)
4. PBS: Phosphate buffer saline

### 2.2. Consumables

1. Parafilm
2. Petri-dishes
3. PAP pen (Abcam ab2601)
4. Poly-L-Lysine Adhesive Microscope Slides (see **Note 1**)

### 2.3. Microscope specifications

1. A range of commercially available atomic force microscopes (AFM) are suitable for studying the mechanical properties of cells by force spectroscopy. The example data presented here was acquired using the following:

a. FlexAFM (Nanosurf AG, Switzerland).
b. mounted on an IX73 (Olympus, Japan) inverted microscope.
2. Measurements were carried out in an air-conditioned laboratory at 21°C.
3. An AFM cantilever with appropriate tip geometry, stiffness and coating material should be chosen.

a. For the present work, PPP-CONTR (Nanosensors, Switzerland) cantilevers with a nominal spring constant of 0.2 N m^1^ and tip radius of 7 nm were used
b. Further details on cantilever selection can be found in **Notes 2-4**.
4. All calibrations and measurements were carried out using Nanosurf C3000 software (Nanosurf AG, Switzerland).

## 3. Methods

### 3.1. Yeast growth and preparation

1. Inoculate 2 ml of YPD with a single yeast colony from a newly-streaked plate and grow overnight at 30°C.
2. For imaging cells on a glass slide, draw a circle by PAP pen to form a hydrophobic barrier for inoculating yeast cells in the next day. Alternatively, place a ring-shape parafilm on the glass slide and heat the glass-slide on a preheated block for 10 seconds to melt the parafilm and form a barrier.
3. If the glass slide is not pre-coated, deposit 100 μl of 0.1% (w/v) poly-L-lysine (PLL) on a glass slide to cover the area and leave the slide in a petri-dish (airdry) for the experiment next day.
4. Dilute the overnight culture in 5 mL of fresh YPD at an OD600 of 0.2 and grow at 30°C for at least one doubling time (>90 minutes but less than 3 doublings).
5. Remove the remaining poly-L-lysine solution on the glass slide and wash it by PBS once, following another wash using YPD medium (or medium suitable for the experiment).
6. Dilute the yeast culture for >100 times in YPD medium and spot 50-100 μl of the diluted culture within the ring area on the glass slide. Keep the slide statically in the room temperature for 30 minutes.
7. Remove the medium gently and wash the area with YPD medium for (at least) twice to remove suspended cells. Add 50-100 μl of YPD medium to keep cells in a constant condition and ready for microscopy.

### 3.2. AFM setup and calibration

1. The procedure for many of the setup steps listed below will be specific to each instrument, and should be carried out following manufacturer’s instructions. In this protocol, a schematic diagram of the experimental setup is shown in Figure 1.
2. Load the selected AFM cantilever into the cantilever holder and attach this to the AFM scan head. When handling probes with a sharp tip, it is recommended to use an electrostatic discharge (ESD) wrist strap and mat to prevent static discharge.
3. Adjust the height and position of the AFM scan head and inverted microscope optics so that the end of the cantilever beam is focused in the field of view.
4. Position the laser beam onto the back of the cantilever, centred laterally and near the tip end, then centre the reflection of the laser on the photodetector.
5. Set up the AFM software in contact mode.
6. Place a clean glass microscope slide on the sample stage of the inverted microscope and pipette a 0.2 ml drop of YPD (or a solution used to keep cells under their natural state; e.g., PBS) onto the centre of the slide.
7. Place the AFM scan head on the stage and lower the cantilever until it is submerged in the droplet. It may help to pipette a small droplet onto the cantilever holder before lowering towards the droplet, to prevent an air bubble being trapped at the interface.
8. Calibrate the cantilever spring constant. Most AFMs have the ability to carry out this calibration using the thermal tuning method (Hutter and Bechhoefer 1993). Where this is not available it is recommended to purchase pre-calibrated probes, as the true value can vary significantly from the nominal value.
9. Approach the cantilever to the surface and calibrate its deflection sensitivity using the glass substrate.
10. Withdraw the cantilever from the surface and remove the glass slide from the stage.

**Figure 1.**
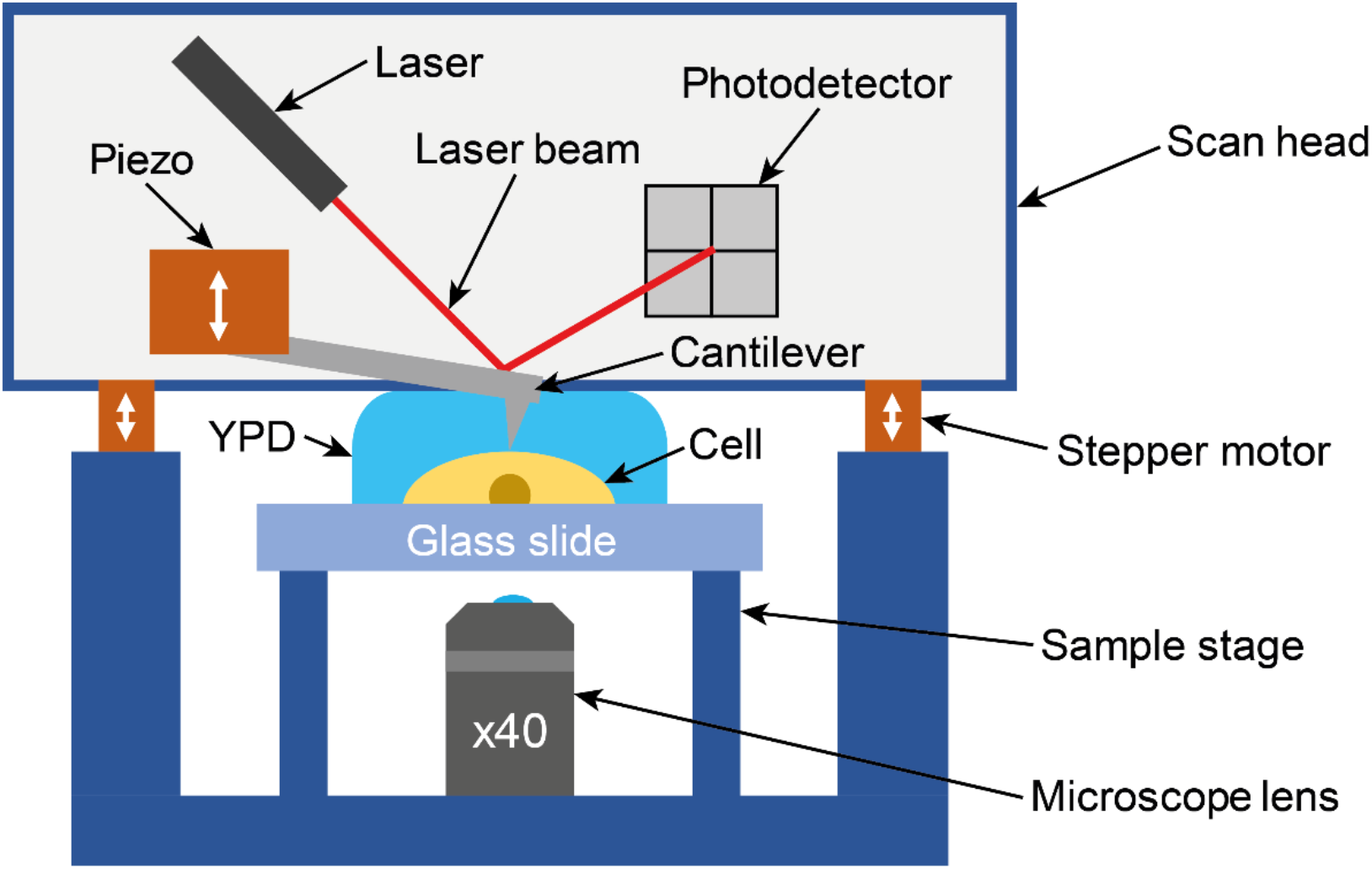
Schematic representation of an Atomic Force Microscope (AFM) instrument mounted on an inverted microscope with glass slide with affixed yeast cell on sample stage.

### 3.3. Force spectroscopy measurements

1. Set up the AFM software for force spectroscopy, selecting the appropriate mode to stop at a maximum force rather than a fixed distance.
2. Input the required parameters (example values are provided below)

a. Maximum force: 1 nN
b. Modulation time: 1 sec
c. Distance range: 5 μm
d. Data points: 1024
e. Grid size: 6 x 6 points
3. To ensure results are reproducible, it is advised to carry our force spectroscopy measurements close to the cell centre. Measurements near the edge of the cell may lead to an increased probability of variation due to the surface gradient of the cell. Either a single measurement at the cell centre, or a grid of measurements over the cell surface may be used. Using a grid allows investigation of localised behaviour over the cell area.
4. Place the poly-L-lysine (PLL) coated glass slide with immobilised yeast cells on the microscope sample stage with a droplet of YPD (based on experimental needs) coating the surface.
5. Lower the cantilever into the droplet.
6. Focus the microscope on the cells.
7. Slowly bring the cantilever towards the surface, stopping when the beam starts to come into focus indicating it is close to the cell surface.
8. Move the sample so the cantilever tip is above the centre of a cell and approach the cantilever to the surface.
9. Start the force spectroscopy measurements.
10. Once the process has been completed, withdraw the cantilever from the cell surface, move the tip to above another cell and carry out repeat measurements.

### 3.4. Data pre-processing

Data analysis can be carried out using a range of commercial or open-source software packages, or using custom written analysis codes in programming languages such as MATLAB or Python (Gavara 2017). The example data here was analysed using a home developed ‘nanoforce’ (v0.3.21) package implemented in Python (v 3.8.1), available from https://pypi.org/project/nanoforce/. The steps used are similar for each platform and are summarised below.

1. Export raw data from the AFM software and import it into a desired analysis package.
2. The output signal from the photodetector measures cantilever deflection as a voltage signal that should be converted to force data using the spring constant and deflection sensitivity calibrated as described in section 3.2. Some AFM software may apply this conversion automatically.
3. The mean value of a selected range of the non-contact portion of the force distance curve is calculated and subtracted from the force data to align the baseline region to 0 nN (as shown in Figure 2b). For some datasets, it may be necessary to instead calculate a linear fit of the non-contact portion and subtract this from the force data, to account for any tilting of the baseline.
4. The contact point at which the tip reaches the sample surface is identified and subtracted from the distance data, so the contact point is at 0 nm (as shown in Figure 2b).
5. Distance data is recorded by the piezoelectric positioner used to control the cantilever height. Due to the physical bending of the cantilever, this may not be able to represent the absolute tip position when in contact with a surface. The true tip position could be calculated from the piezo position, cantilever deflection, and deflection sensitivity.
6. Figure 2a presents a force-distance curve acquired on the PLL substrate as benchmark. This was recorded to distinguish between the cell surface and substrate.
7. Figure 2b shows a force-distance curve measured on the surface of a cell identified by the optical microscopy.
8. Figure 2a and 2b show distinctively different characteristics: in the approaching component of the force curve (blue lines), a much deeper indentation can be observed on the cell than the PLL substrate, and there is a considerable hysteresis on the cell surface when withdrawing the cantilever.
9. A range of information can be quantified from an AFM force-distance curve, including the following which are labelled in Figure 2b:

a. Adhesion force (N) – maximum force recorded on the withdraw curve when separating the AFM tip from the surface
b. Work of adhesion (J) – area between the baseline and the withdraw curve when separating the AFM tip from the surface
c. Young’s modulus (Pa) – calculated from the slope of the approach curve in the indentation region

**Figure 2.**
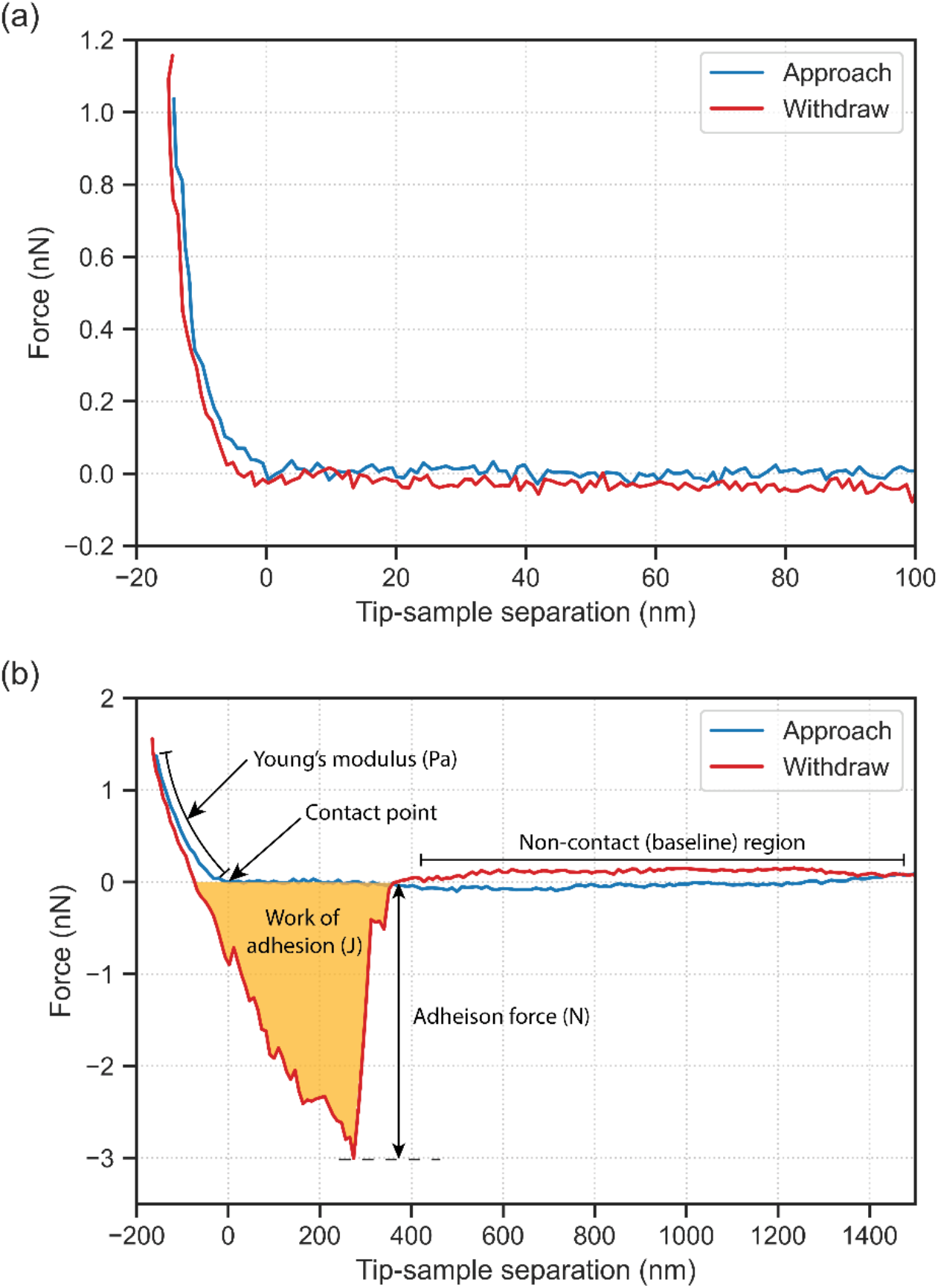
Representative AFM force-distance curves for (a) PLL substrate used to immobilise cells, used to exclude measurements where cantilever did not contact cell surface, and (b) cell surface with key parameters annotated.

### 3.5. Contact mechanics analysis

1. A range of contact models reported in the literature can be used to calculate Young’s modulus from a force-distance curve. An appropriate model should be selected depending on the geometries of the surface (i.e. the cell) and the indenter (i.e. the AFM tip) (Han and Chen 2021).
2. The cell can be assumed a flat surface if its size dimension is much greater than that of the indenter.
3. For a spherical indenter on a flat surface (Figure 3a), the force-distance curve can be fitted to the Hertz model (Eq. 1) (Johnson 1985).

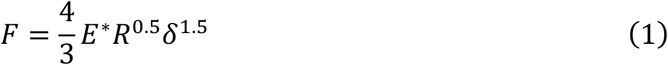

where *F* is force (N), *R* is tip radius (m), *δ* is indentation depth (m) and *E*\* is reduced modulus (Pa), which is given by Eq. 2.

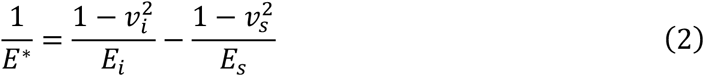

where *E* is Young’s modulus (Pa) and *v* is Poisson ratio, with subscript *i* representing the indenter (AFM cantilever) and s the sample. Assuming a rigid indenter (i.e. Eq. 3) with a Young’s modulus much greater than that of the sample, simplifies to Eq. 4.

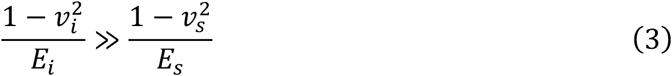

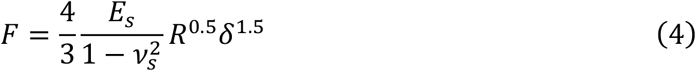
4. For a conical indenter on a flat surface (Figure 3b) the force-distance curve can be fitted to the Sneddon model (Eq. 5) (Chen 2014).

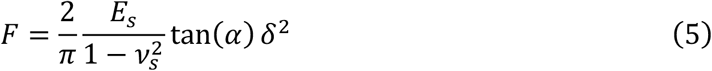

where *α* is the half angle of the indenter.
5. Hertz (Eq. 4) and Sneddon (Eq. 5) model fits of the approaching force-distance curve (Figure 2b) are shown in Figure 4a alongside the raw data points.
6. Data can be fitted to each model by taking a linear fit of the force versus distance terms (i.e. *F* vs. *δ*^1.5^ for the Hertz model and *F* vs. *δ*^2^ for the Sneddon model), using an ordinary least squared linear regression model.
7. The gradient of the calculated linear fit can be substituted into the equation for the appropriate model, along with the Poisson ratio and tip radius or tip half-angle, to calculate the Young’s modulus of the sample.
8. The Sneddon model fitting in Figure 4**Error! Reference source not found.**a is a much better representation of the raw data than the Hertz model fitting. This is expected as the AFM cantilever used to record this data has primarily conical shape, excluding a small spherical region at the tip with a radius of less than 7 nm.
9. Figure 4b shows a pointwise analysis of the Young’s modulus at each raw datapoint, along with the value predicted by the Sneddon model fitting shown in Figure 4a. The model proves an accurate representation of the raw data points at indentation depths of greater than 40 nm.
10. The divergence from the model at low indentation depths is likely caused by the small spherical region at the tip of the cantilever. Once the indentation depth is much greater than this tip radius, the cantilever can be assumed cylindrical, for which the Sneddon model fits well.
11. Literature investigating the adhesion and Young’s modulus of yeast cells has observed variation in measured values across the cell surface (Çolak et al. 2020; Formosa et al. 2013).
12. Figure 5 shows the variation of values measured over the surface of 3 representative cells.

**Figure 3.**
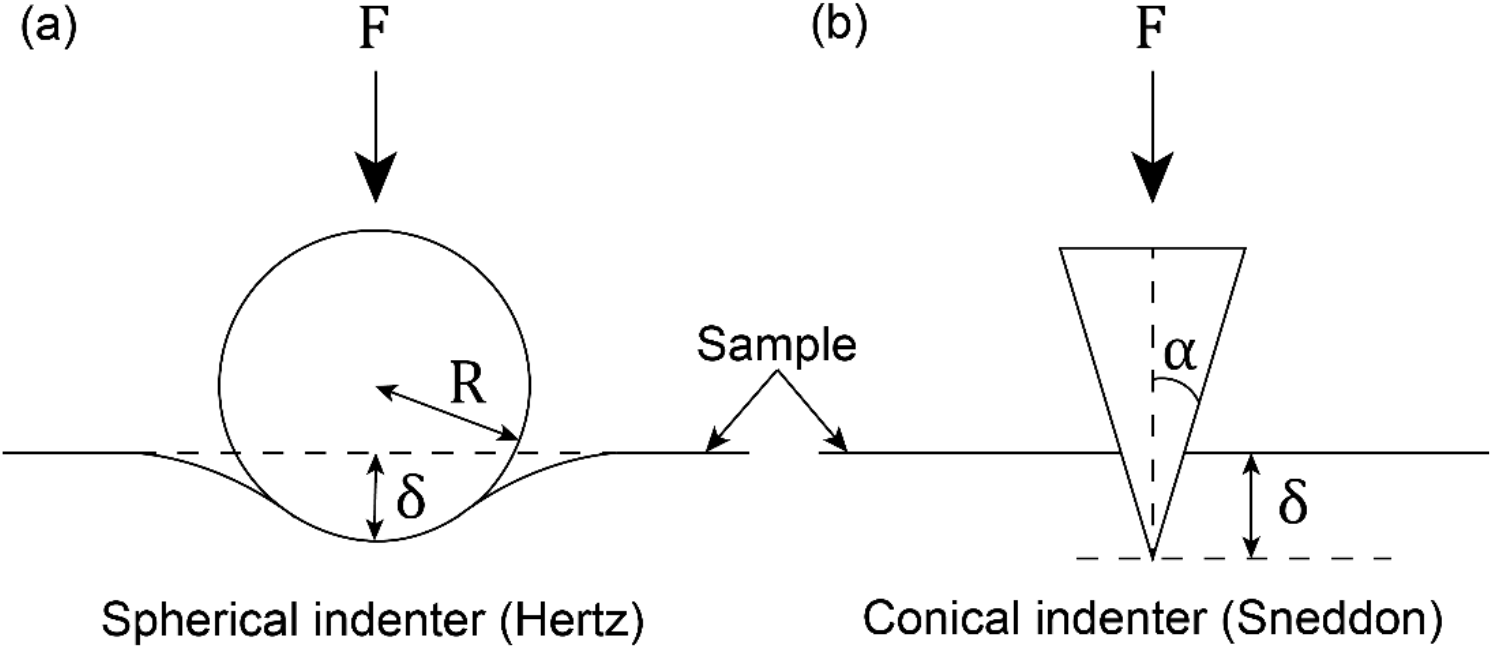
Schematic representation diagrams of the indentation of a flat sample by (a) a spherical indenter and (b) a conical indenter with the relevant parameters required to fit force-distance data shown for the Hertz and Sneddon models respectively.

**Figure 4.**
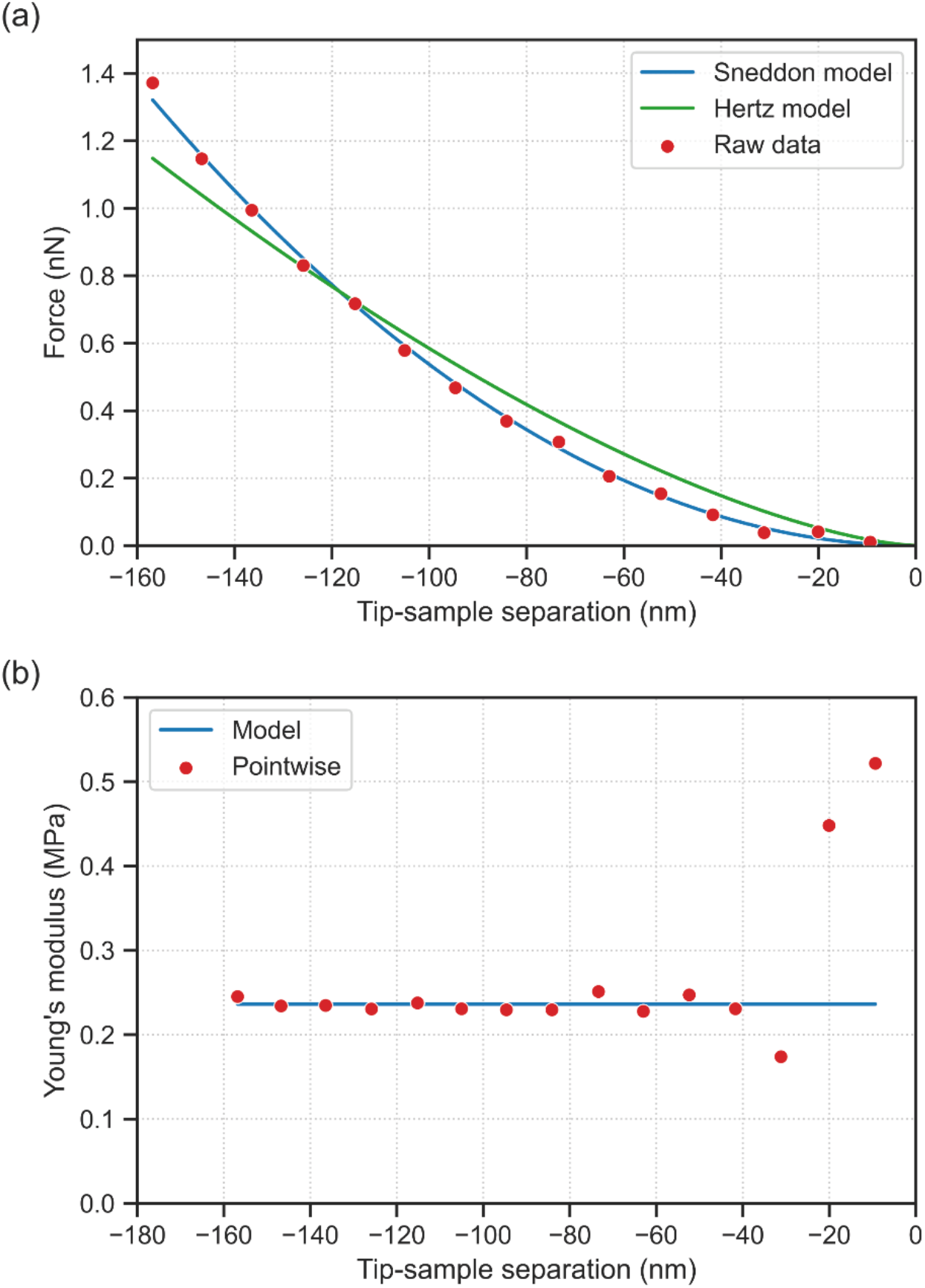
(a) Raw force-distance data for indentation portion of AFM measurements, along with Sneddon model and Hertz model fits of this data, and (b) comparison of Young’s modulus calculated by the Sneddon model using the fit shown in (a), along with a pointwise calculation for each raw data point.

**Figure 5.**
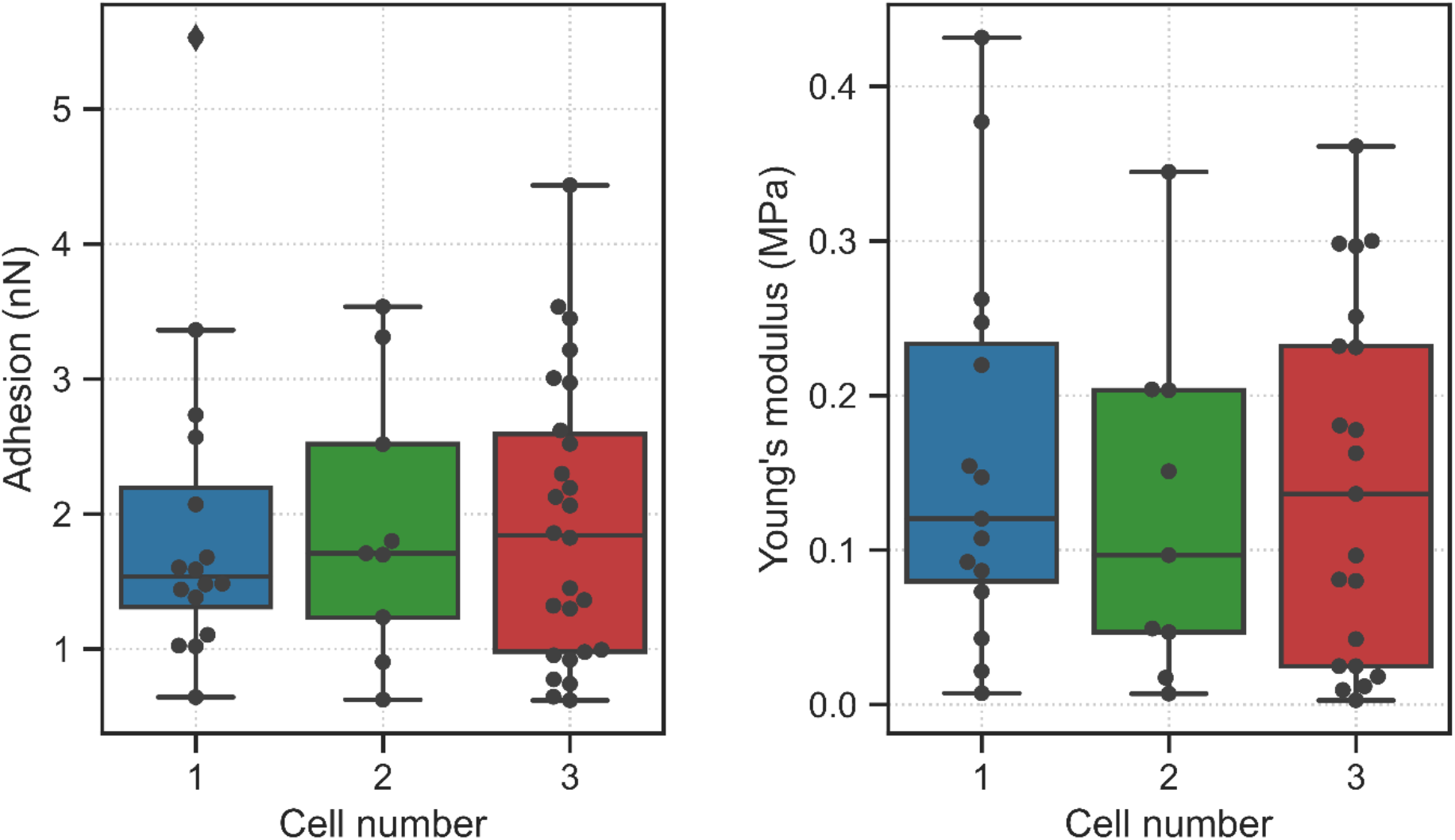
Box plots showing the variation of adhesion force and Young’s modulus for three representative cells.

## 4. Notes

1. Yeast cell immobilization using an adhesive (e.g., poly-L-lysine and concanavalin A) is recommended, because yeast cell may move away while the cantilever makes contact. Cell entrapment using microfabricated chambers/microwells may alter the mechanical properties in their natural state since the space confinement could apply an artificial force to the cell.
2. A cantilever should be chosen with a spring constant that allows large cantilever deflection to improve the resolution of the force-distance curve while being stiff enough to allow sufficient indentation into the cell and ensure the cantilever detaches from the cell surface during the withdraw curve. Typically, a spring constant between 0.01 and 0.6 N/m is appropriate for cell studies (Gavara 2017).
3. Cantilever tip geometry is an important consideration for studying the nanomechanical properties of cells. In general, a spherical tip with a 2.5-10 μm radius is most appropriate for soft and fragile cellular structures, while a conical tip may be used for stiffer features. The sharp tip of a conical indenter enables determination of more localised surface properties, while a spherical tip gives information related to the whole-cell behaviour (Chen 2014).
4. When aligning the AFM tip with the cell surface by optical microscopy it may be advantageous to use a cantilever with the tip aligned with the end of the cantilever beam for ease of alignment, such as OPUS 3XC-NA (Mikromasch, USA) or ARROW-CONTR (NanoWorld, Switzerland).

## 5. Acknowledgement

The work is supported by the Royal Society Research Grant RGS\R2\202400 to H-J. T.; Engineering and Physical Science Research Council Grants EP/V029762/1 to Z.J.Z. and EP/R511845/1 to C.R.J.

